# Effects of auditory reliability and ambiguous visual stimuli on auditory spatial discrimination

**DOI:** 10.1101/2020.06.11.140491

**Authors:** Madeline S. Cappelloni, Sabyasachi Shivkumar, Ralf M. Haefner, Ross K. Maddox

**Affiliations:** Biomedical Engineering, University of Rochester, Rochester, NY 14627, USA; Del Monte Institute for Neuroscience, University of Rochester, Rochester, NY 14627, USA; Center for Visual Science, University of Rochester, Rochester, NY 14627, USA; Brain and Cognitive Sciences, University of Rochester, Rochester, NY 14627, USA; Neuroscience, University of Rochester, Rochester, NY 14627, USA

## Abstract

The brain combines information from multiple sensory modalities to interpret the environment. Multisensory integration is often modeled by ideal Bayesian causal inference, a model proposing that perceptual decisions arise from a statistical weighting of information from each sensory modality based on its reliability and relevance to the observer’s task. However, ideal Bayesian causal inference fails to describe human behavior in a simultaneous auditory spatial discrimination task in which spatially aligned visual stimuli improve performance despite providing no information about the correct response. This work tests the hypothesis that humans weight auditory and visual information in this task based on their relative reliabilities, even though the visual stimuli are task-uninformative, carrying no information about the correct response, and should be given zero weight. Listeners perform an auditory spatial discrimination task with relative reliabilities modulated by the stimulus durations. By comparing conditions in which task-uninformative visual stimuli are spatially aligned with auditory stimuli or centrally located (control condition), listeners are shown to have a larger multisensory effect when their auditory thresholds are worse. Even in cases in which visual stimuli are not task-informative, the brain combines sensory information that is scene-relevant, especially when the task is difficult due to unreliable auditory information.

## I. INTRODUCTION

When we navigate our surroundings, we encounter sensory information from multiple modalities. Combining complementary information across sensory modalities often helps us construct a more accurate percept of the world. In contrast, combining conflicting or irrelevant sensory information can lead to perceptual errors. In order to optimize perceptual accuracy, the brain must determine whether to combine information across sensory modalities, and if so, how to weigh each sensory modality. Formally, the notion of reliability weighting is described by Bayesian models of cue combination—forced integration (Ernst and Banks, 2002) and more recently causal inference (Körding et al., 2007). In these models, each cue is treated as a measurement of the stimulus with a Gaussian distribution of the likelihood of the stimulus based on that measurement. The multisensory measurement is then a combination of unisensory measurements weighted by the inverse of their relative variances, such that a narrower likelihood distribution will have more influence on the combined percept. Importantly, the causal inference model adds another layer of inference to this model, in which the degree of cue integration depends on the probability that both measurements actually arose from the same event in the world (Körding et al., 2007).

Bayesian models of multisensory integration are typically tested in tasks in which the subject can use information from multiple modalities to determine the correct response; for example, an audiovisual localization task in which the subject is asked where a noise and light occurred (Körding et al., 2007). Under good visual conditions, this task gives rise to the “ventriloquist effect”, a bias of auditory location towards the visual stimulus (Howard and Templeton, 1966). However, when the visual stimulus gets blurrier and harder to localize relative to the auditory stimulus, the apparent visual bias weakens or even manifests as an auditory bias of perceived visual location (Alais and Burr, 2004). This demonstrates that the ventriloquist effect is truly a bias of both visual and auditory stimuli towards each other with the magnitude of the bias determined by the relative reliability of each modality. Importantly, in this and other tasks described by the model, there is only one stimulus in each sensory modality, both of which are informative about the correct response. In this scenario, it is optimal for the brain to use multisensory integration to improve its judgment and behavioral performance.

Previously, we extended the classical work by increasing the number of visual and auditory cues and found a multisensory effect of visual stimuli on auditory spatial processing (Cappelloni et al., 2019) even when those visual cue did *not* contain any task-relevant information. We asked listeners to perform a concurrent auditory spatial discrimination task in which random visual stimuli were either spatially aligned with two symmetrically separated auditory stimuli or both collocated in the center of the screen, and found a performance benefit when the visual stimuli were spatially aligned. This audiovisual effect goes against the traditional conception of multisensory integration as a mechanism for the optimal combination of information from the environment (Ernst and Bülthoff, 2004). The benefit provided by the spatially aligned visual stimuli is also not explained by an ideal Bayesian observer (whose response should be invariant to the locations of the task-uninformative visual stimuli), and is counterintuitive in that the visual stimuli do not help to determine the correct response and must instead benefit the listener through process not part of the ideal Bayesian observer model.

Here we test the hypothesis that the brain weighs auditory and visual stimuli by their relative reliabilities even in the case where the visual stimuli do not provide any information about the correct response and would be ignored by an ideal observer. We modulated the reliability of the auditory stimuli by changing their duration, with longer auditory stimuli being more reliable. We found that the benefit provided by the visual stimuli is larger where subjects had poor auditory thresholds. Our results replicate those of our previous study (Cappelloni et al., 2019) and further investigate the ways in which scene-relevant but task-uninformative stimuli can shape perception providing constraints for future theoretical models.

## II. METHODS

### A. Participants

Participants (16 female, 4 male; ages ranging between 18 and 31, mean 21.5 +/- 3 years) with normal hearing (thresholds 20 dB HL or better at 500-8000 Hz) and normal vision (self-reported) gave written informed consent. They were compensated for the full duration of time spent in the lab. Research was performed in accordance with protocol approved by the University of Rochester Research Subjects Review Board.

### B. Stimuli

Auditory stimuli were pink noise tokens and harmonic tone complexes with matching spectral envelopes bandlimited to 220–4000 Hz. Stimuli were generated and localized by HRTFs from the CIPIC library using interpolation from python’s expyfun library as in (Cappelloni et al., 2019), with the notable difference that we generated the pink noise tokens and harmonic tone complexes to be three durations, 100 ms, 300 ms, and 1 s. Auditory stimuli were presented at a 24414 Hz sampling frequency and 65 dB SPL level from TDT hardware (Tucker Davis Technologies, Alachua, FL) over ER-2 insert earphones (Etymotic Research, Elk Grove Village, IL).

Visual stimuli were regular polygons of per-trial random number of sizes and color. They were inscribed within a 1.5° diameter circle. Colors were chosen to have uniform saturation and luminance, with the two stimuli in each trial having opposite hue as in (Cappelloni et al., 2019). Visual stimuli had the same onset and offset times as the auditory stimuli and thus matched their duration. To prevent overlap they were presented in alternating frames (Blaser et al., 2000) on a monitor with a 144 Hz refresh rate.

### C. Task

Each trial began when the subject fixated on a white dot in the center of the screen, confirmed with an eye tracking system (EyeLink 1000, SR Research). Then all four auditory and visual stimuli were presented concurrently for the duration of the trial (100 ms, 300 ms, or 1000 ms). After stimulus presentation, subjects were asked to respond with what side the tone was on by pressing a button. There were two visual conditions: one in which the visual stimuli were spatially aligned with the auditory stimuli and one in which the visual stimuli were collocated in the center of the screen.

We presented trials according to weighted one up one down adaptive tracks that adjusted the separation of the two sounds (Kaernbach, 1991). Separations were adjusted on a log scale such that separation increased by a factor of 2 when the participant responded incorrectly and decreased by a factor of **2**^**1/3**^ when they responded correctly. Each track had 130 trials and began at a starting separation of 10° azimuth. For each track, we randomized the number of trials with the tone on the left and right. There were six tracks, three durations by two visual conditions, that were randomly interleaved.

### D. Analysis

In order to obtain 75% thresholds we averaged the level at each reversal (skipping the first six reversals). Threshold improvement is defined as the difference between the separation thresholds of the two visual conditions (central – matched). We resampled the reversals with replacement to determine the significance of each threshold improvement (positive or negative respectively – less than 2.5% or greater than 97.5% of resampled threshold improvements less than zero). We performed linear regression on the central threshold vs. threshold improvement data and computed 95% confidence intervals using the Python seaborn package (Michael Waskom et al., 2017). We also fit a linear mixed effects model to the data using Python’s statsmodels package (Seabold and Perktold, 2010). We fit the thresholds with a random intercept model such that each subject is assigned an intercept to control for between subject variance. We considered categorical visual condition, duration, and interaction of visual condition and duration as fixed effects in the model.

## III. RESULTS

Subjects improved their task performance, indicated by a decrease in threshold, asymptotically in both visual conditions as the duration of the auditory stimuli increased; however, there was considerable variation in subject performance. Only 11 of 20 subjects were able to perform the auditory discrimination task at the shortest duration such that we could calculate a separation threshold (Figure 1A). Subjects in this group had a large decrease in threshold between 100 ms and 300 ms, but did not improve further for 1000 ms stimuli. For the remainder of the subjects who had thresholds too big to calculate in either or both 100 ms conditions, they improved their threshold between 300 ms and 1000 ms. In a linear mixed effect model of all subjects combined, only duration (p=9.5×10^−9^) had a significant effect on threshold. Visual condition and the interaction of visual condition and duration did not have a significant effect on threshold.

**Fig. 1.**
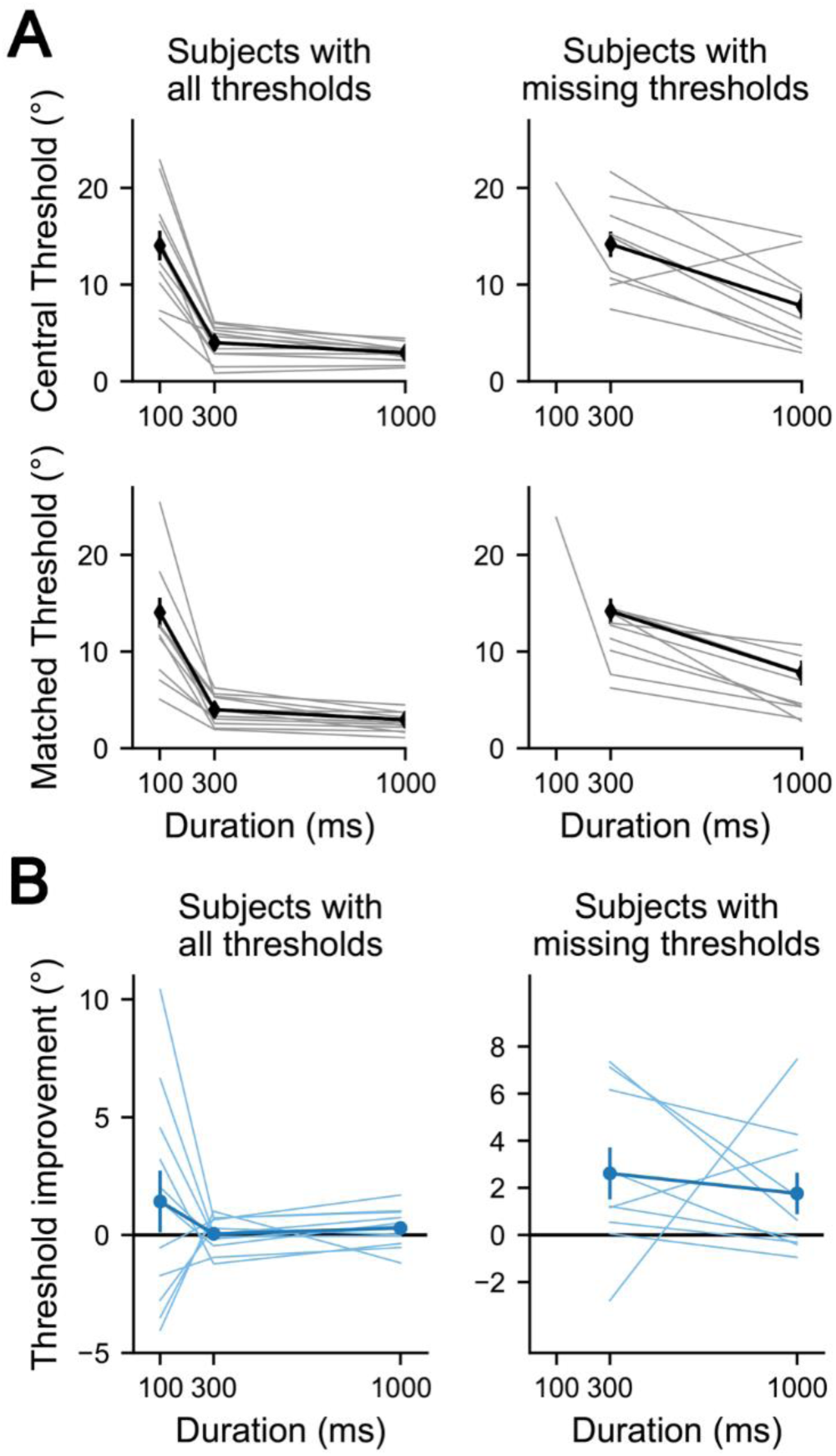
(Color Online). A. Thresholds for each duration in the central visual condition (top) and matched visual condition (bottom). Many subjects had missing thresholds (too large to measure accurately) in one or both visual conditions at 100 ms and are plotted in the right column (n=9) while the remainder are plotted on the left (n=11). B. Improvements in threshold for the two groups of subjects: those who could perform the task at all durations (left), those had one or both thresholds missing at 100 ms (right).

We defined “threshold improvement” as the difference between the central and matched visual conditions and used it to measure the size of the visual benefit (Figure 1B). Differences in individual auditory spatial processing ability indicate that auditory reliability was not uniform within a given duration condition.

In order to compensate for individual differences we used the separation threshold in the central condition as a measure of auditory reliability for each duration condition (Figure 2). Larger thresholds indicated that the task was more difficult and the individual’s auditory reliability was likely worse. Pooling measurements across durations, we found a linear relationship between the threshold improvement and central threshold (Figure 2D, r=0.53, p=5×10^−5^). A similar trend is shown when plotting data from our previous study (Cappelloni et al., 2019) (only using 300 ms stimuli) on the same axes (Figure 2E, r=0.76, p=10^−4^).

**Fig. 2.**
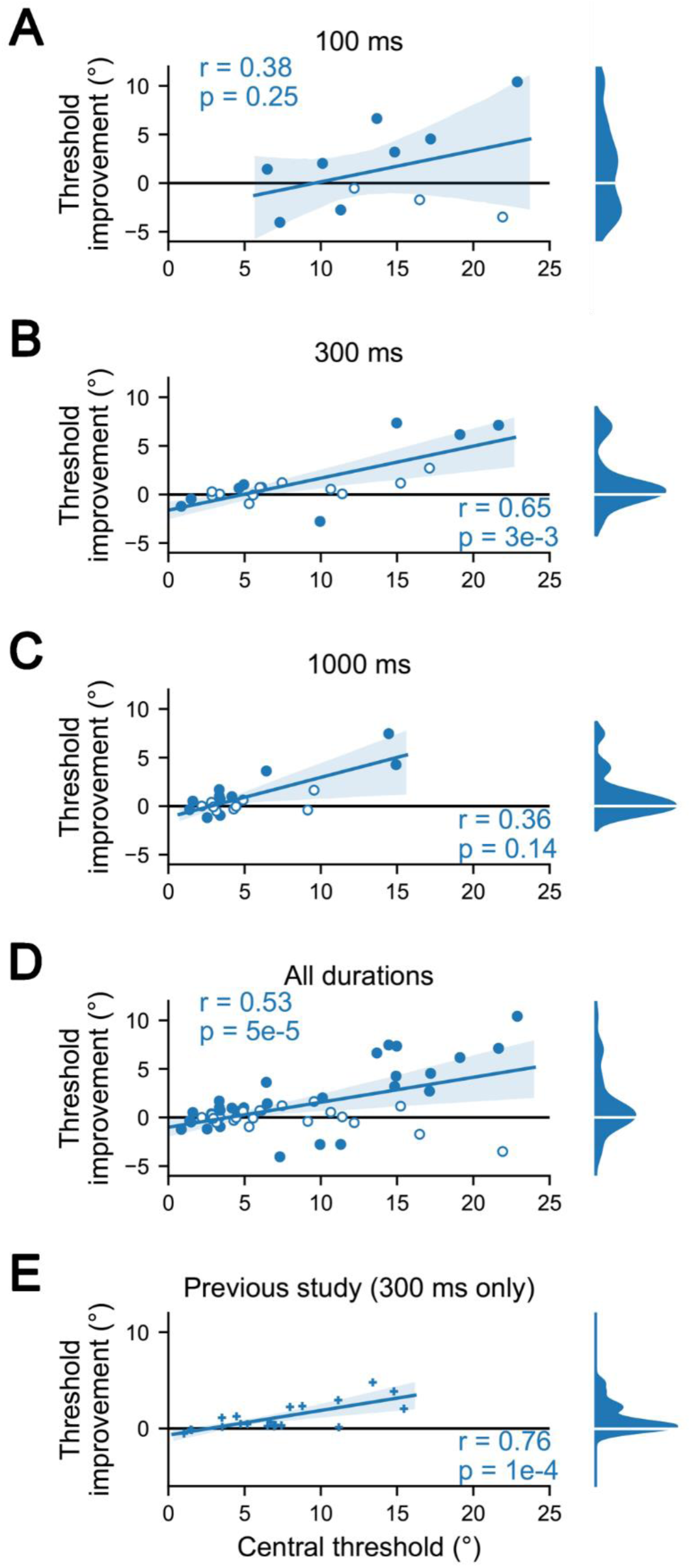
(Color Online). Linear regression of threshold improvements against central threshold. Solid markers indicated significant differences between the two visual conditions based on within subject variation (**α = 0. 05**, uncorrected). Open markers indicate no significant effect of visual stimuli. Also shown are marginal kernel density estimates (excluding tails beyond which there is less than 0.5% of the mass). A-C. Separate regressions for each duration condition. D. Regression for threshold improvements regardless of duration. E. Regression of data from our previous study (Cappelloni et al., 2019).

## IV. DISCUSSION

We found that performance in the auditory task correlated with the benefit subjects receive from task-uninformative visual stimuli. Listeners experienced the most benefit when the auditory task was difficult for them (large central threshold). In contrast, individuals who did well in the auditory task (small central threshold) experienced no benefit or even a slight decrement.

It should be noted that as the central threshold gets larger, the stimuli become more peripheral where spatial acuity is worse (Hafter and Maio, 1998; Maddox et al., 2014; Middlebrooks and Onsan, 2012; Mills, 1958), compounding listeners’ worse auditory reliability. A similar decrease may also occur in visual location reliability. Extending the duration improves listener’s thresholds in both visual conditions, which can be explained by an improvement in the auditory reliability, suggesting that our duration manipulation does roughly correlate with reliability.

This work replicates our previous finding that task uninformative but spatially aligned stimuli benefit auditory spatial discrimination. In the previous study, the visual stimuli preceded auditory stimuli by 100 ms, whereas in this study, their onsets were all concurrent. Additionally, we previously used sigmoidal fits to determine threshold instead of adaptive tracks. This suggests that the visual benefit is robust to small audiovisual asynchronies and changes in probabilistic distribution of stimuli across space (adaptive tracks will lead to more non-uniform priors).

Although the visual benefit was larger where subjects showed worse auditory performance, and duration had a significant effect on task difficulty, we did not see a significant effect of changing the duration on visual benefit. This is mainly due to the wide range of auditory spatial processing abilities among the subjects. Because of differences in auditory spatial processing, the effect of duration on visual benefit was inconsistent across subjects. Subjects could be divided roughly into two groups with two different patterns of thresholds, those who could reliably perform the task at 100 ms and those who could not. The former group improved their performance significantly when the duration was extended to 300 ms, but did not further improve when the stimuli were 1000 ms, suggesting that they reached ceiling performance at 300 ms. In contrast, the latter group improved significantly when the stimulus duration extended from 300 ms to 1000 ms and were not at ceiling performance at 300 ms. In both this study and our original experiment (Cappelloni et al., 2019), which only included the 300 ms duration condition, we observed a wide range of auditory thresholds. In addition to simple variability across subjects, some thresholds were missing data points because the monitor on which visual stimuli were displayed could not extend far enough to accurately measure large thresholds. These missing data may have allowed us to better fit a model that could show an effect of changing the stimulus duration on visual benefit if such an effect existed. Without considering effects on the scale of individual subjects, for which we did not have enough data, we could not establish a causal relationship between changing stimulus duration and the visual benefit even though correlations suggest one may exist.

It is possible that auditory reliability is the underlying factor driving the relationship between auditory performance and visual benefit, even though our data did not show a clear relationship between duration and visual benefit. If auditory reliability does modulate the effect of task-uninformative visual stimuli, following the spirit of Bayesian causal inference, it would further violate the central assumption that the brain will only integrate information that is relevant to the task. This would point to a broader notion of multisensory perception in which stimuli are integrated based on their reliability in representing the sensory scene, rather than the reliability of information they provide regarding a specific task. Further work is needed to describe the boundaries of what information is integrated in a scene. It is important that such work go beyond traditional paradigms to those that can reveal differences of scene-relevant vs task-informative cues

## V. CONCLUSION

We show that listeners gain a larger benefit from task-uninformative visual stimuli in an auditory spatial discrimination task when the auditory task is difficult. Our results are consistent with, but do not confirm, the notion that reliability weighting as described in Bayesian models may occur even when visual stimuli do not carry information about the correct decision in the task. We believe two next steps would clarify the findings of this paper. “Small-n” design in which few subjects are recruited to complete many trials would allow us to understand perception on the level of individuals, rather than generalizing across a diverse population (Smith and Little, 2018). Secondly, we call for an exploration of more complex paradigms with multiple multimodal cues caused by potentially multiple events in the world that provide new and stronger tests of existing models.

## ACKNOWLEDGMENTS

The authors wish to acknowledge Sara Fiscella for assisting with data collection.

Research reported in this publication was supported by the National Institute on Deafness and Other Communication Disorders of the National Institutes of Health under award number R00DC014288.

## REFERENCES

Alais, D., and Burr, D. (2004). “The Ventriloquist Effect Results from Near-Optimal Bimodal Integration,” Curr. Biol., 14, 257–262. doi: 10.1016/j.cub.2004.01.029

Blaser, E., Pylyshyn, Z. W., and Holcombe, A. O. (2000). “Tracking an object through feature space,” Nature, 408, 196-.

Cappelloni, M. S., Shivkumar, S., Haefner, R. M., and Maddox, R. K. (2019). “Task-uninformative visual stimuli improve auditory spatial discrimination in humans but not the ideal observer,” PLOS ONE, 14, e0215417. doi: 10.1371/journal.pone.0215417

Ernst, M. O., and Banks, M. S. (2002). “Humans integrate visual and haptic information in a statistically optimal fashion,” Nature, 415, 429–433. doi: 10.1038/415429a

Ernst, M. O., and Bülthoff, H. H. (2004). “Merging the senses into a robust percept,” Trends Cogn. Sci., 8, 162–169. doi: 10.1016/j.tics.2004.02.002

Hafter, E. R., and Maio, J. D. (1998). “Difference thresholds for interaural delay,” J. Acoust. Soc. Am., 57, 181. doi: 10.1121/1.380412

Howard, I. P., and Templeton, W. B. (1966). Human spatial orientation, Human spatial orientation, John Wiley & Sons, Oxford, England, 533 pages.

Kaernbach, C. (1991). “Simple adaptive testing with the weighted updown method,” Percept. Psychophys.,.

Körding, K. P., Beierholm, U., Ma, W. J., Quartz, S., Tenenbaum, J. B., and Shams, L. (2007). “Causal Inference in Multisensory Perception,” PLOS ONE, 2, e943. doi: 10.1371/journal.pone.0000943

Maddox, R. K., Pospisil, D. A., Stecker, G. C., and Lee, A. K. C. (2014). “Directing Eye Gaze Enhances Auditory Spatial Cue Discrimination,” Curr. Biol., 24, 748–752. doi: 10.1016/j.cub.2014.02.021

Michael Waskom, Olga Botvinnik, Drew O’Kane, Paul Hobson, Saulius Lukauskas, David C Gemperline, Tom Augspurger, et al. (2017). mwaskom/seaborn: v0.8.1 (September 2017), Zenodo. doi: 10.5281/zenodo.883859

Middlebrooks, J. C., and Onsan, Z. A. (2012). “Stream segregation with high spatial acuity,” J. Acoust. Soc. Am., 132, 3896–3911. doi: 10.1121/1.4764879

Mills, A. W. (1958). “On the Minimum Audible Angle,” J. Acoust. Soc. Am., 30, 237–246. doi: 10.1121/1.1909553

Seabold, S., and Perktold, J. (2010). “Statsmodels: Econometric and Statistical Modeling with Python,” Austin, Texas, 92–96. Presented at the Python in Science Conference. doi: 10.25080/Majora-92bf1922-011

Smith, P. L., and Little, D. R. (2018). “Small is beautiful: In defense of the small-N design,” Psychon. Bull. Rev.,, doi: 10.3758/s13423-018-1451-8. doi:10.3758/s13423-018-1451-8

